# Weakly-Supervised Tumor Purity Prediction From Frozen H&E Stained Slides

**DOI:** 10.1101/2021.11.09.467901

**Authors:** Matthew Brendel, Vanesa Getseva, Majd Al Assaad, Michael Sigouros, Alexandros Sigaras, Troy Kane, Pegah Khosravi, Juan Miguel Mosquera, Olivier Elemento, Iman Hajirasouliha

**Affiliations:** Institute for Computational Biomedicine, Department of Physiology and Biophysics, Weill Cornell Medicine, New York, NY, USA; Department of Theoretical and Applied Science, Ramapo College of New Jersey, Mahwah, NJ, USA; Englander Institute for Precision Medicine, Weill Cornell Medicine, New York, NY, USA; Department of Pathology and Laboratory Medicine, Weill Cornell Medicine, New York, NY, USA; Computational Oncology, Department of Epidemiology and Biostatistics, Memorial Sloan Kettering Cancer Center, New York 10021, USA; The Meyer Cancer Center, Weill Cornell Medicine, New York, NY, USA

## Abstract

Estimating tumor purity is especially important in the age of precision medicine. Purity estimates have been shown to be critical for correction of tumor sequencing results, and higher purity samples allow for more accurate interpretations from next-generation sequencing results. In addition, tumor purity has been shown to be correlated with survival outcomes for several diseases. Molecular-based purity estimates using computational approaches require sequencing of tumors, which is both time-consuming and expensive. Here we propose an approach, weakly-supervised purity (wsPurity), which can accurately quantify tumor purity within a slide, using multiple and different types of cancer. This approach allows for a flexible analysis of tumors from whole slide imaging (WSI) of histology hematoxylin and eosin (H&E) slides. Our model predicts tumor type with high accuracy (greater than 80% on an independent test cohort), and tumor purity at a higher accuracy compared to a comparable fully-supervised approach (0.1335 MAE on an independent test cohort). In addition to tumor purity prediction, our approach can identify high resolution tumor regions within a slide, to enrich tumor cell selection for downstream analyses. This model could also be used in a clinical setting, to stratify tumors into high and low tumor purity, using different thresholds, in a cancer-dependent manner, depending on what purity levels correlate with worse disease outcomes. In addition, this approach could be used in clinical practice to select the best tissue block for sequencing. Overall, this approach can be used in several different ways to analyze WSIs of tumor H&E sections.

## Introduction

In recent years, there has been an increase in tumor DNA sequencing from cancer patients, to identify tumor driving somatic mutations. However, samples that are used for sequencing consist of not only cancer cells, but also include normal immune cells, stromal cells and blood vessels, whose DNA also gets sequenced. The purity of tumor samples impacts sequencing results, making it harder to detect mutations, especially subclonal events due to dilution of tumor DNA by normal DNA. Determining tumor purity is therefore critical before sequencing. Moreover, purity in some cases can provide prognostic information. In glioma and colon cancer, low tumor purity is associated with worse survival outcomes^1,2^. Therefore, tumor purity estimates can help correct sequencing outputs and predict disease outcomes.

Traditionally, tumor purity has been estimated by pathologists based on review of hematoxylin and eosin (H&E) stained slides. Several methods, including ABSOLUTE^3^, ESTIMATE^4^, as well as a consensus purity estimate (CPE)^5^, have aimed to determine tumor purity through computational analysis of sequencing results. Molecularly derived purity scores, especially those relying on DNA, are attractive because they are thought to be highly accurate and do not need any manual review or inspection.

However, sequencing and associated analysis are expensive processes. On the other hand, pathologist-derived purity scores may have limited accuracy because they may have substantial inter-pathologist variability; indeed previous literature has shown that pathology-estimated tumor purity often fail to correlate well with sequencing-derived purity^6^. Consensus-based approached have been one solution proposed for this problem, but they require either multiple pathologists or multiple methodologies for molecular approaches, increasing monetary and time costs^6^. Thus, other approaches to quantify tumor purity are necessary.

H&E-stained slides are cheap and quick to produce from tumor samples and have been the gold standard for cancer diagnosis by trained pathologists. The utilization of artificial intelligence (AI) methods, in particular, deep learning for analyzing histopathology images has significantly increased over the past few years. Notable deep learning methods were developed to identify the presence of tumor in a tissue, segment regions of interest, and classify molecularly-derived subtypes^7–16^. There have been two main approaches to handling these types of deep learning analyses, fully-supervised and weakly-supervised. A fully-supervised approach requires each image to have a label associated with it to train a model. In fully supervised approaches, such as the data that was presented for the CAMELYON16 and CAMELYON17 datasets^17^, expert pathologists will review all slides used for training and testing an algorithm and manually annotate the slides to represent classes or segmented regions of interest. Due to computational constraints, entire gigapixel sized images, with several thousand pixels in height and width, cannot be directly used as input into standard deep learning techniques, although sophisticated approaches have been proposed to make these large images compatible^18^. Instead, the consensus approach to date has been to split the slides into small patches or tiles (hundreds of pixels for width and height), that can be efficiently fed into these models. In this scenario, annotated regions are necessary as there may be small subsections of the tissue, at the size of a single patch, that have no tumor tissue within it. It is possible to train a model using slide level labels for patches, but this will result in noisy labels, as some non-tumor patches within a given tumor slide will be given the wrong label. The problem with a fully-supervised approach, in which manual annotations of regions is needed, is that it requires a significant amount of time from domain experts to label enough slides to be used to properly train deep learning models.

Therefore, recent works have focused on using weakly-supervised approaches.

In a weakly-supervised approach, instead of manually annotating the entire slide, region by region or patch by patch, the slide is given a single overall label, and deep learning models use this information to identify regions of interest or use this to classify the disease state of the slide as a whole. This allows a much quicker annotation process, as the entire slide can be given a label and the heterogeneity within the slide can be accounted for. The main premise of most weakly-supervised approaches requires pooling the features from image patches, under the multiple instance learning (MIL) framework^19^. In this scenario, a “bag” represents the entity in which the classification occurs, and “instances” represent heterogeneous elements that all relate to the bag entity. In the case of the standard assumption for MIL, if at least one instance within the bag is positive, the whole bag can be considered positive^19^. This concept was applied by Campanella et al.^8^, where the authors used > 15,000 cancer patients with several years’ worth of histological slides from the Memorial Sloan Kettering Cancer Center (MSKCC) slide repository in addition to external patient slides to predict the presence of cancer within a slide. The pooling operation used in this approach was max pooling. In this case, a slide is positive if at least one patch is predicted as positive (positive would be the presence of tumor). The authors provided a proof-of-concept study that clinical-grade performance (described as an AUC of greater than 0.98 for all cancer types for cancer detection) was achieved using this approach, and that specific malignant regions within slides were identified^8^. As mentioned by Lu et al, this requires a lot of data, since very few patches from each slide are used for the classification; in fact, only one patch per slide is used during backpropagation for the max pooling operator^15^. Another pooling operation that can be used is average pooling under the collective assumption of the MIL algorithm. In this regard, all of the instances within a bag, have an equal influence on the bag class^19^. A seminal work was introduced in 2018, called Attention-MIL, which introduced a new pooling approach and is a modification of this collective assumption^20^. The principle of this approach is that a neural network is used within the model to learn a good aggregating function. In this sense, it is learning a weighted sum of all the instances within a bag, to make the prediction of interest. The different weights can then be interpreted as different types of data that have varying impacts on model prediction. This approach has been used to find slides that contain cancer and identify cancer subtypes^15^. It has also been used to automatically identify the hormonal status in breast cancer from H&E slides^21^, stage of cancer^22^, and tissue origin of metastatic lesions^23^.

In this study, we propose a weakly-supervised approach to calculating tumor purity from whole slide sections utilizing an attention-based, multi-task, multiple instance deep learning model, we call wsPurity, for whole-slide purity detection. Most of the previous work done using weakly-supervised methods have provided labels to perform binary or multi-class classification, indicating 1 for the correct class label and 0 for the rest. Previous work has incorporated tumor purity in the prediction task^24^, however this was performed in a fully-supervised manner. In this work, we propose that incorporating tumor purity into the classification task will allow for the proper identification of tumor regions within the histological slide, and improved performance over this fully-supervised approach.

We use an Attention-MIL setup to learn a weight feature for the distribution of patches within a slide and a feature representation that can accurately predict tissue type as well as tumor purity level. We adapt a similar deep learning pipeline to the multi-task multiple instance learning approach used in Lu et al.^23^, but the fully-connected layers, specific tasks, and loss functions applied differs from their approach. Framing the purity score prediction as an ordinal classification problem, wsPurity predicts tumor purity within 9 different bins ranging from 0 to 1, where 0 is no tumor and 1 is complete tumor. We use a pathologist derived consensus purity score developed from previous literature as the ground-truth to train our model based on TCGA database slides^25^. There are several unique benefits for our model, which include 1) We can accurately identify tumor purity in a tissue slide and compare our results to previously developed models, 2) We can classify tumors into low and high purity at several different thresholds that can be cancer type specific, and 3) we can identify potential tumor regions that can be isolated and used to enrich the tumor sample for improved sequencing.

## Results

A total of 5,850 slides, comprised of six different cancer types, including adrenal adenocarcinoma (ACC), lung squamous cell carcinoma and lung adenocarcinoma (LUSC & LUAD respectively), invasive breast carcinoma (BRCA), head and neck squamous cell carcinoma (HNSC), prostate adenocarcinoma (PRAD), and ovarian serous cystadenocarcinoma (OV), were used to train, validate, and test our proposed deep learning model (70% | 15% | 15% respectively). The most frequent tumor types were invasive breast carcinoma (BRCA) and lung cancer (LUAD & LUSC), a combination of lung adenocarcinoma and lung squamous cell carcinoma (Supplementary Table 1). The model was tested using two separate cohorts of patients. The TCGA database slides were used for model training and validation (4,063 and 921 slides respectively), and a held-out test set (866 slides) from the TCGA database was used to evaluate model performance (TCGA cohort). In addition, we used a Weill Cornell Medicine (WCM) cohort of 54 de-identified H&E slides for evaluating model performance and generalizability (WCM independent cohort). We also evaluate how differences in the amount of cancer tissue available from the TCGA database and the ratio of cancer tissue slides to normal tissue slides in the TCGA database affect model performance on a per cancer basis.

There is variability in the purity distribution between different cancer types. Figure 2 shows the distribution of the held-out TCGA test set purity distribution stratified by tumor type. Here we can see that tumor purity distributions are cancer-type specific, for example OV is skewed towards higher tumor purity, whereas PRAD is skewed to lower tumor purity, when excluding normal tissues. In addition, pathology provided tumor purities overall are skewed towards higher values, 0.7 (0.35-0.85) - median (IQR), for all slides used in this study. In addition, if the normal tissue slides are removed, the values for median (IQR) shift higher to 0.8 (0.67 – 0.89).

The overall workflow of our model can be seen in Figure 1A. WSIs first get split into a set of tiles, and tiles are then filtered to remove out of focus regions and regions that have little to no tissue present. These patches get passed through a deep learning model, where each original image patch is transformed into a feature representation. These patches are combined using a weighted sum of each feature vector, using an attention mechanism, to obtain two final representations that can be used for two downstream tasks, prediction of tumor purity and prediction of tissue type. Figure 1B shows the schematic for the proposed wsPurity deep learning model. In particular, we used a previously developed Resnet-IBN^26^ network that has been shown to improve generalizability of deep learning models, especially when there is the presence of color variation, which is common for H&E stained slides. The building block of the Renset-IBN model is shown in Figure 1C.

**Figure 1:**
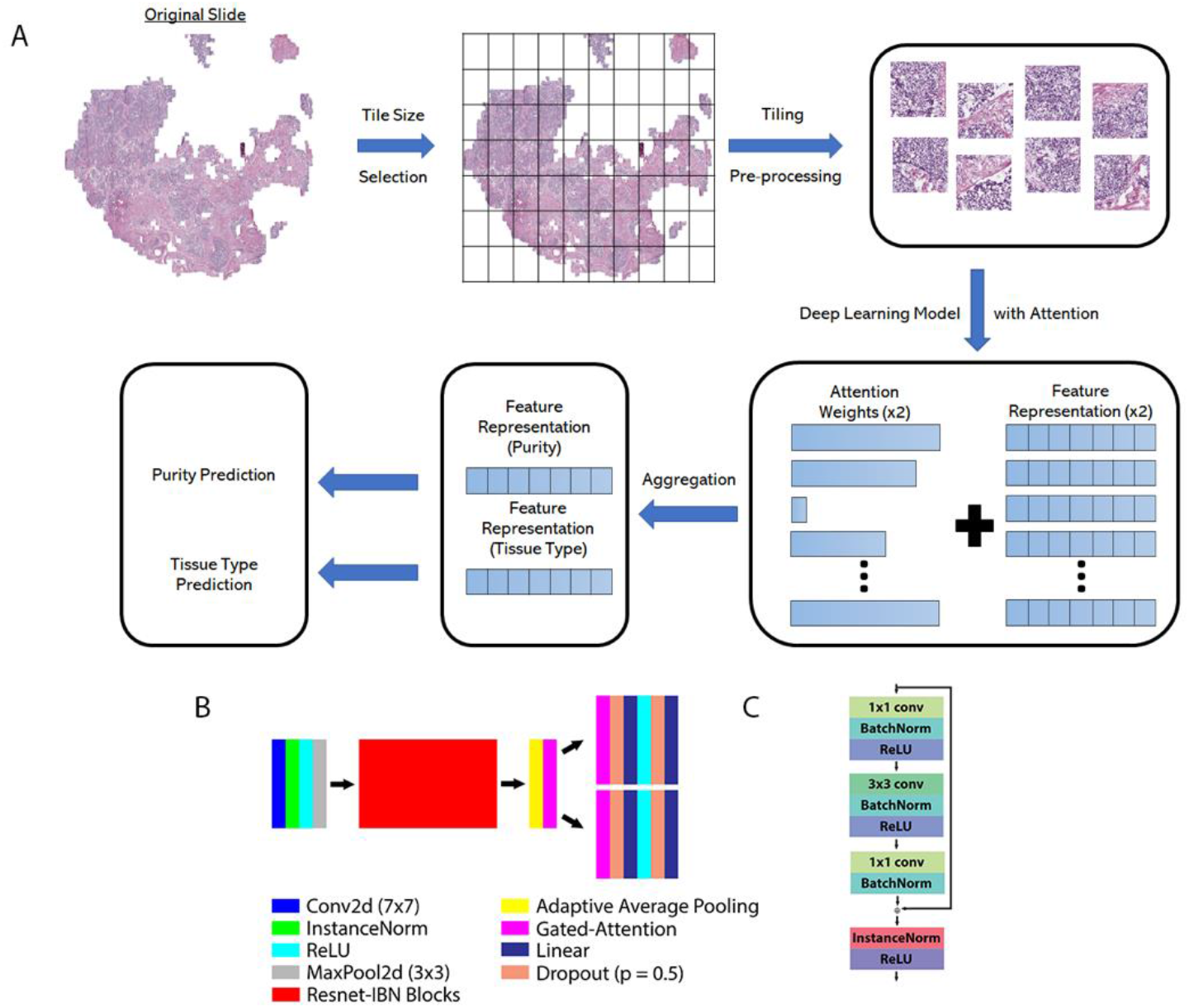
(A) Workflow of wsPurity. To get a slide output, the original svs file is tiled, passed through a deep learning model with an attention mechanism to combine information from all tiles to perform two tasks, tissue type prediction and tumor purity prediction. (B) Schematic of the multi-attention multi-task MIL approach. The model uses the structure of Resnet-34-IBN, which is a modified Resnet model using InstanceNorm. A gated attention mechanism generates two feature representations, which pass through a set of linear and dropout layers for the final predictions. (C) Schematic of a residual block from the Resnet-34-IBN-b model (Right - Adapted from Pan et al.^26^)

### Cancer Type Prediction

We first aimed to be able to predict cancer type for a given tumor. As shown in Table 1, model performance for tissue type prediction was high, and was consistent between the validation and test set – indicating that our model is generalizable to unseen TCGA slides. The average accuracy among the six cancer types is 97.8% (average of six cancer type accuracies). The tissue type that performed the worst was lung cancer at 95%. When analyzing the feature embedding generated from the model, or features generated from bags of 120 patches, we clearly see separations between the six different tissue types.

**Table 1.**
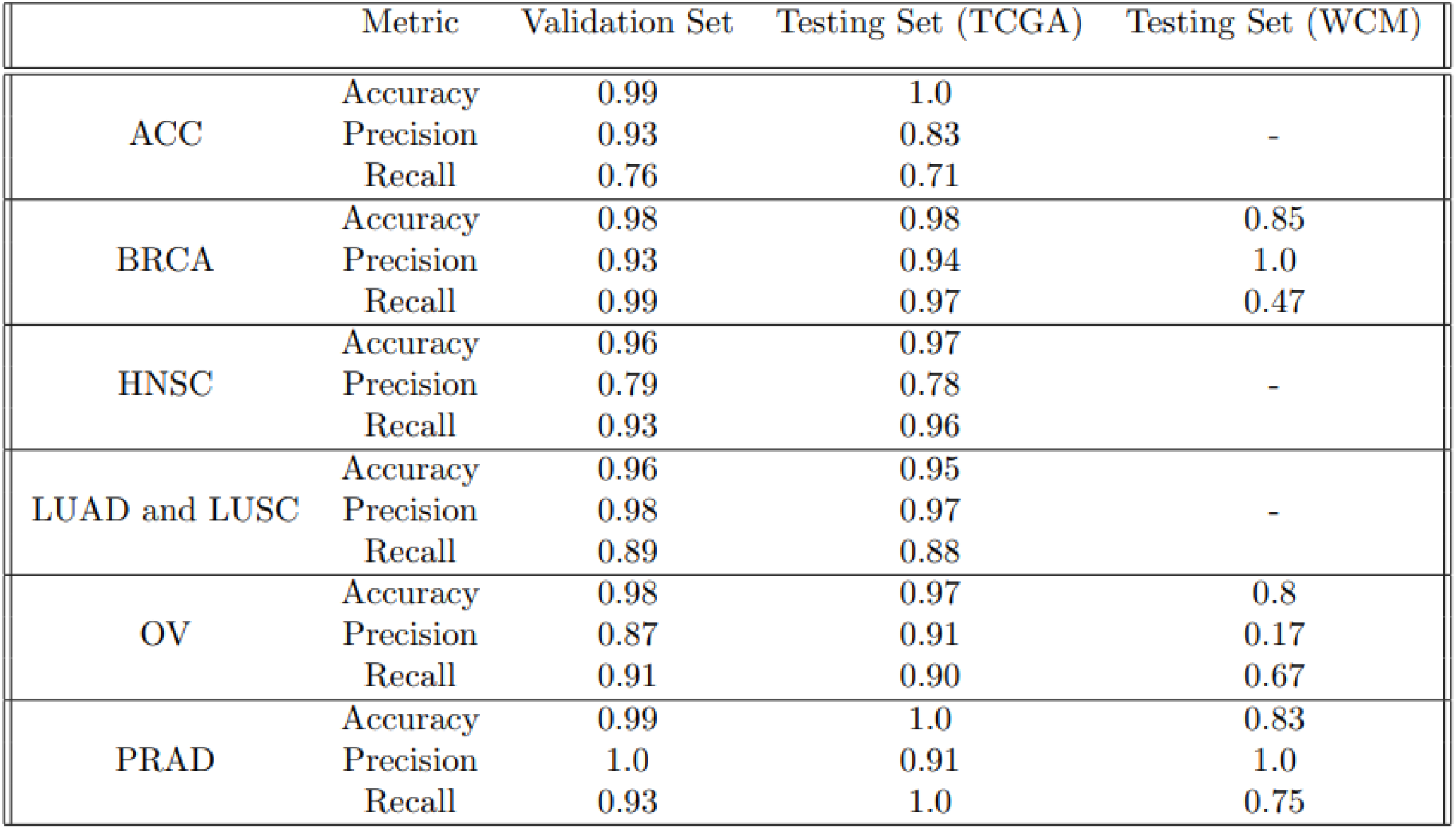
Reported values of accuracy, precision and recall for the tissue type prediction for the validation set, test set (TCGA cohort), and test set (WCM independent cohort)

Interestingly, the features from HNSC and LUAD/LUSC have some mixing with one another based on the tSNE plot (Figure 4). BRCA and PRAD, and OV generally have unique feature representations. We see high model performance for our WCM independent cohort of patients (accuracy greater than 0.8 for all cancers). However, we see a slight drop compared to the TCGA cohort, which could be related to tissue quality or tissue preprocessing differences between cohorts, although further investigation is necessary. The recall values for BRCA and precision value for OV may be skewed due to very low sample size (15 and 3 for BRCA and OV respectively), compared to PRAD (36 slides or ∼ 2/3 of total test set), which has more similar performance to the TCGA cohort of slides (Supplementary Figure 2).

### Tumor Purity Prediction

In addition to evaluating performance metrics for cancer type, we quantified the performance of tumor purity prediction. We compared our model against the fully-supervised approach previously published^24^, as well as compared how our model performed in the held-out TCGA cohort compared to the WCM cohort. We identified 701 slides from our test set that overlapped with previously published work in the Fu et al. paper^25^. To assess how well our model performed, we compared both the mean squared error (MSE) and mean absolute error (MAE) of our model for both comparisons. Firstly, we see that our weakly-supervised approach performs better than previous published work, with a MSE value of 0.0441 vs 0.1538, and we see that the same trend holds for MAE (0.1659 vs 0.2967). In addition, we see that our model also generalizes well to our WCM independent test cohort. The performance of tumor purity prediction on the WCM, although significantly smaller in size, is comparable to that of the TCGA held-out data (0.0279 - MSE and 0.1335 MAE).

In addition, to validate this model in determining clinically relevant subclasses of tumor purity (high vs. low tumor purity), we generated receiver operating characteristic (ROC) curves based on three separate thresholds (Figure 4). The first threshold is for tumor vs. normal tissue prediction. Since we do not have any tumor purity less than 0.09, the 10% threshold can be used to consider the classification between these two groups. What we see is that for the majority of tumor types, the AUC is greater than 0.98. The lowest performing tissue type is HNSC, which may be due to the lower proportion of normal tissue to tumor tissue distribution (∼1:20 normal to tumor ratio). This is much lower than all other cancers besides the most infrequent class, ACC, by a significant amount (the next smallest was PRAD at ∼ 1:5 ratio). This trend follows for the two tumor purity prediction thresholds (60% and 70% thresholds) as well. The tumor type with the best performance in terms of the area under the curve (AUC) of the tumor purity prediction is ACC. This could be due to the very biased tumor purity scores (all values above 80% purity). Overall, model performance decreases when moving the tumor purity threshold higher (Figure 2). The worst performing model was the HNSC tumors (worst AUC: 0.78, 70% threshold), whereas we see high performance (AUC > 0.8) for ovarian cancer, lung cancer and adrenal cancer, and breast cancer for all the classification thresholds.

**Figure 2:**
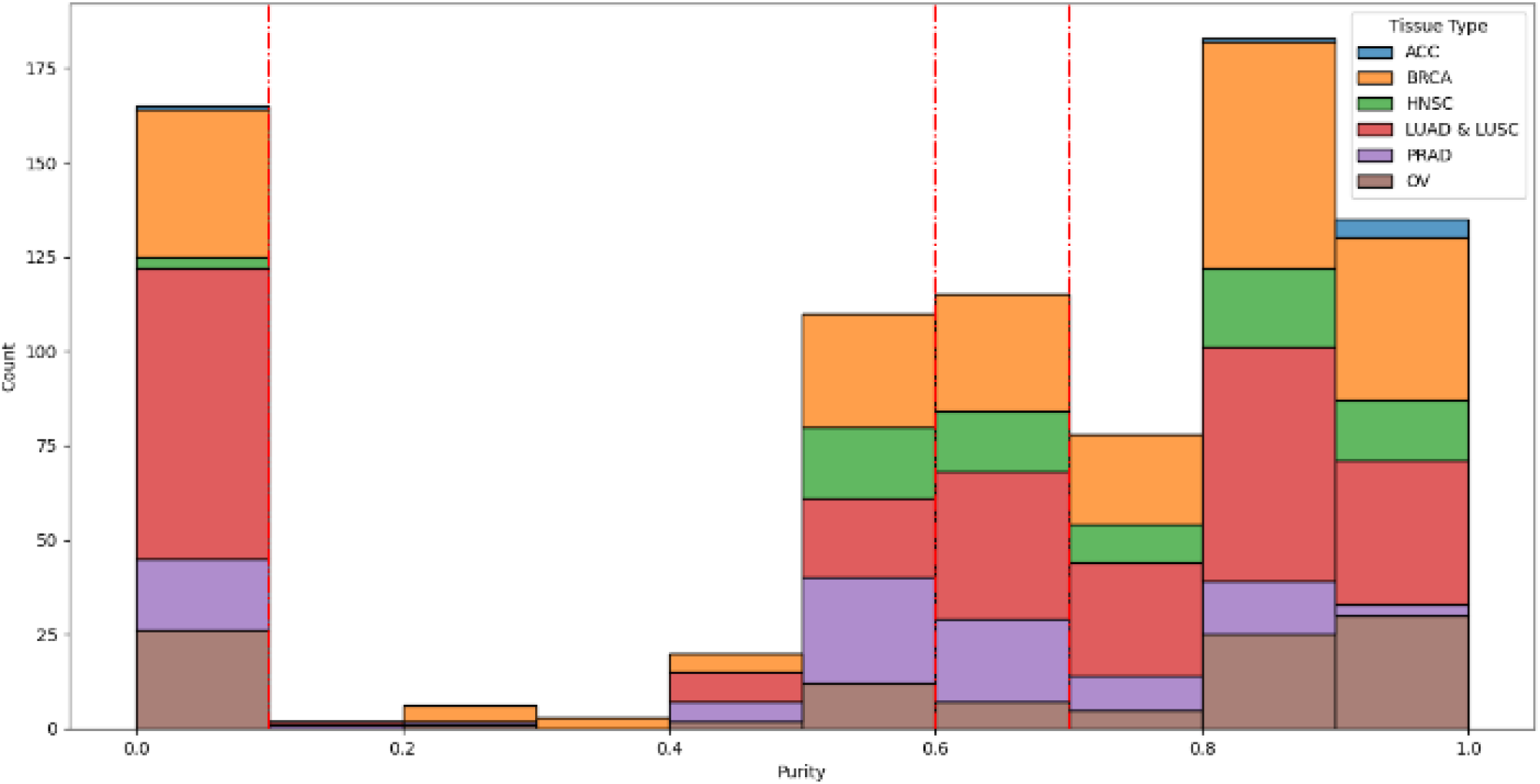
Representative test set distribution for tumor purity data stratified by tissue type. Red lines show the thresholds used for identifying low vs high tumor purity.

When analyzing the feature embeddings for the purity scores, we see that the predicted purity values follow a uniform pattern from highest to lowest. Since these embeddings are on the batch or “bag” level, they may correspond to only a subset of the tissue slice, if the number of tissue patches within a slide exceeded 120, due to computational constraints. Therefore, what we see is that in the extreme cases (i.e. pure tumor or no tumor), the embeddings match well with the predicted tumor type. When the tumor is a mixture of normal and abnormal tissue, we see that there is a combination of many different tumor purities across the slide (Figure 4C, Figure 4D). This is significant, as when analyzing tumor samples, it is important to identify the purest sample for downstream sequencing analyses. We do see some normal tissue classified as tumor based on the embeddings. As seen in the ROC curves (Figure 3), there are errors in tumor vs. normal prediction. The normal tissues however that are predicted as tumor, are predicted as low tumor purity, as seen by the t-distributed stochastic neighbor embedding (tSNE) plots in Figure 4.

**Figure 3:**
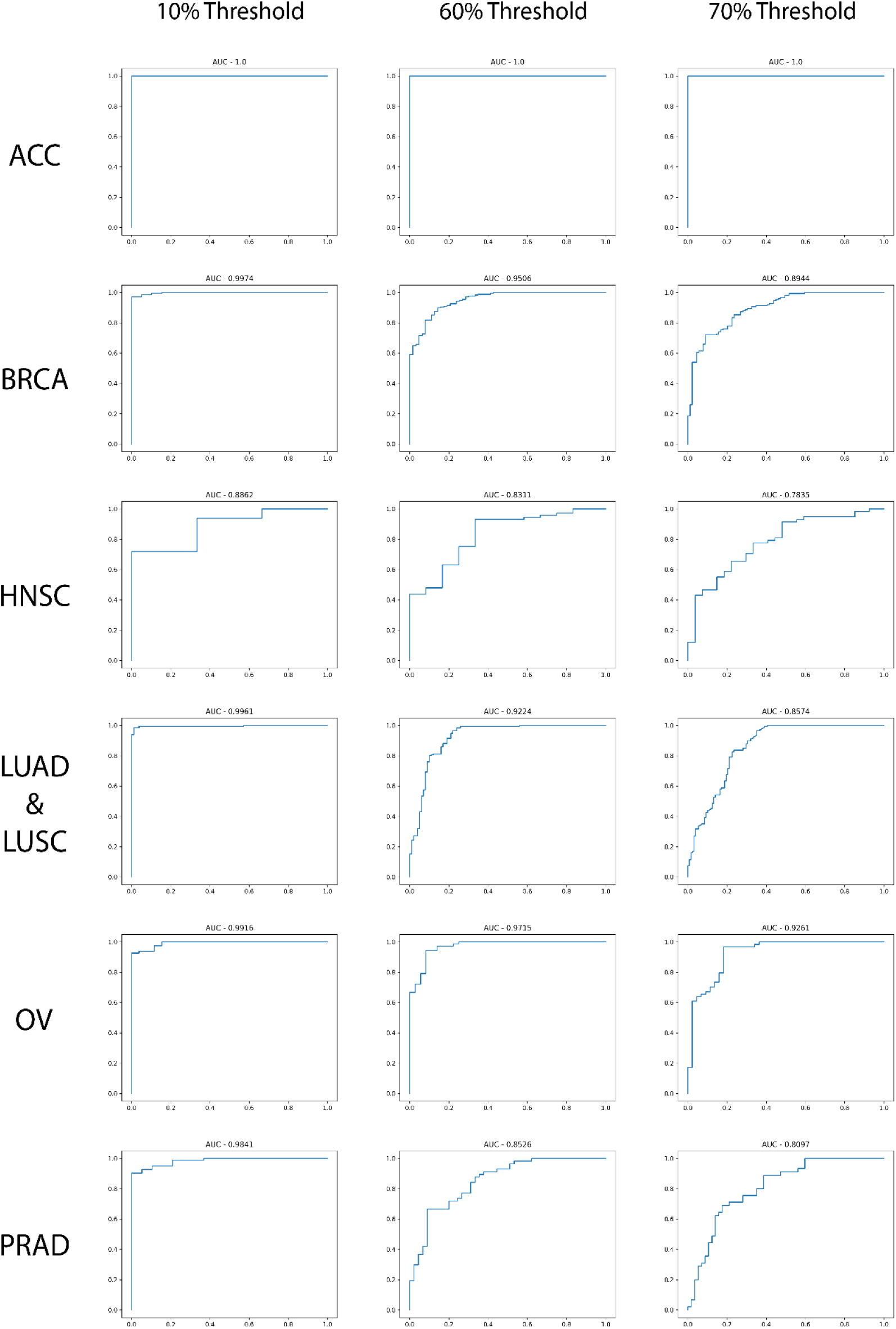
ROC Curves of high vs low tumor purity. We set the thresholds at 10% tumor purity, 60% tumor purity, and 70% tumor purity to identify model performance comparing normal vs. tumor tissue.

**Figure 4:**
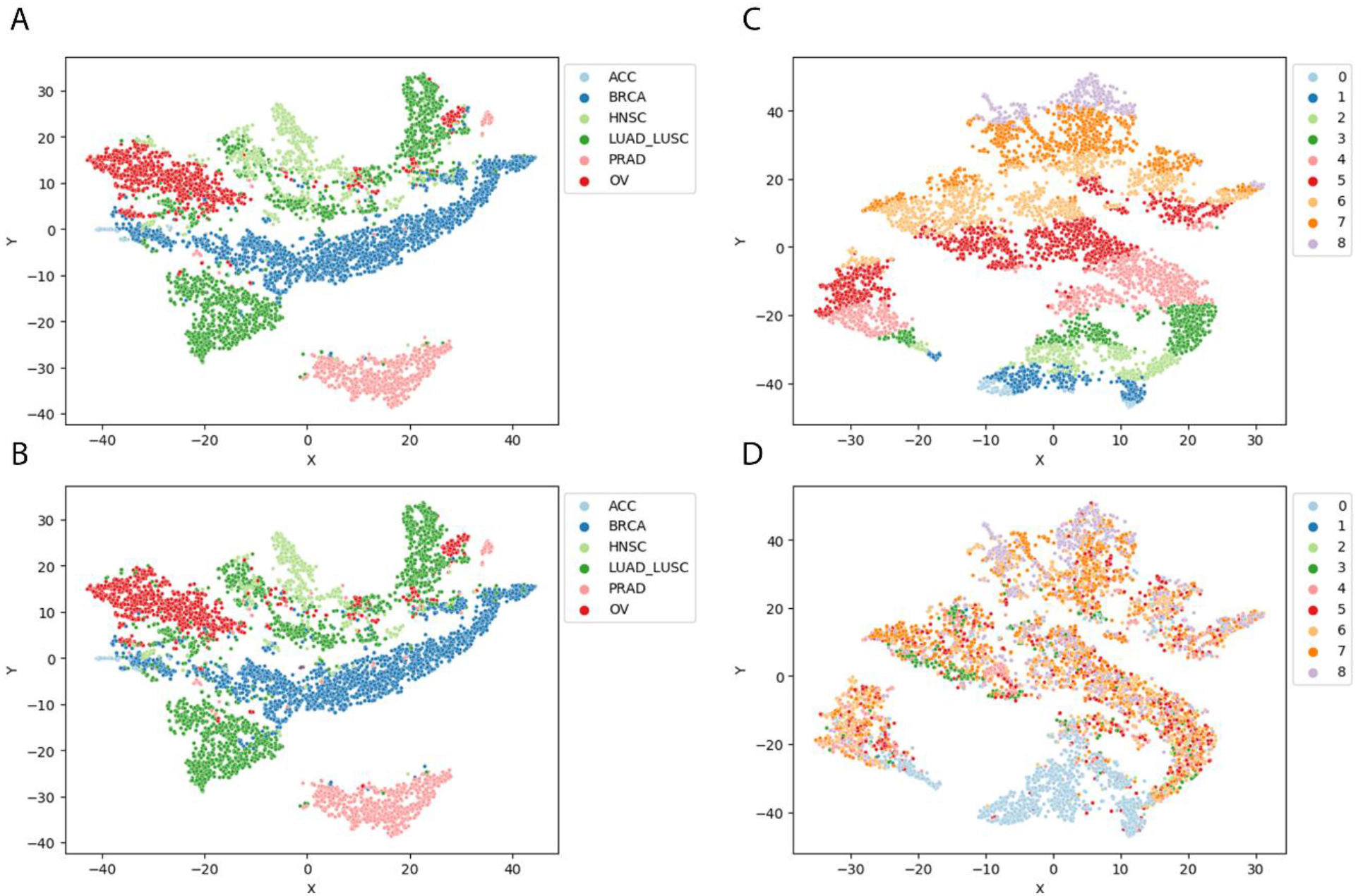
(A-B) tSNE Plots using the feature embedding from the tissue type prediction (compared true vs. predicted, respectively) (C-D) tSNE Plots using the feature embedding from the tumor purity score prediction (compared true vs. predicted, respecti

### Tumor Visualization and Purity Distribution

To assist clinicians and basic scientists, the regions of high tumor purity need to be visualized. In this model, we can identify different regions within the tissue, such as tumor and non-tumor regions (Figure 5). We do this by relying on the attention weights generated from the deep learning model and the output from the purity prediction. Due to the multi-task attention approach, we have two weights per patch per slide, one for tissue type prediction and one for tumor purity prediction. We will focus solely on the tumor purity weights, and compare these regions with pathologist-derived labels of these tumor regions.

**Figure 5:**
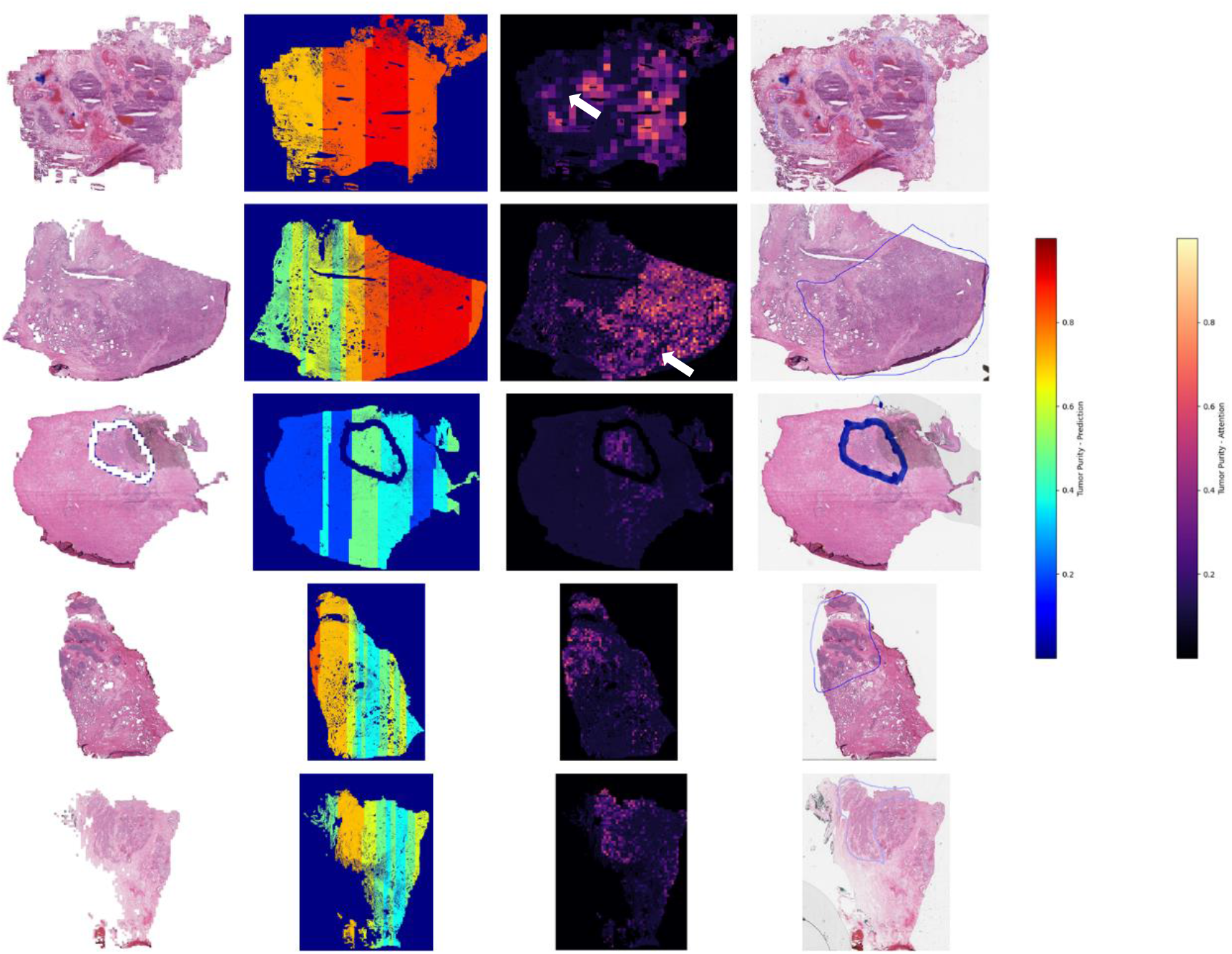
Analysis of the tumor purity prediction based on WCM independent cohort slides. (Left) A view of the overall tissue architecture. (Middle-Left) The distribution of the tumor purity predictions within the entire slide. (A-Middle-Right) The distribution of the attention weights for tumor purity prediction. (A – Right) Pathologist-derived annotations for the tumor region within the slide. White arrows represent regions with low tumor purity identified within pathologist derived regions, showing higher resolution detail for our approach compared to pathologists.

Tissue sections are *a priori* split into batches of 120 tiles, similar to what has been done in previous multi-task learning methods, as computationally all tiles cannot be analyzed simultaneously^8^. We use the *x* and *y* coordinates from each tile, to determine the geographical relationship between tiles. We perform density-based spatial clustering (DBSCAN) on these coordinates, to identify unique tissue sections within the given tissue slide, as multiple sections can be placed on a given slide. We next identify each cluster and sort the *x* coordinates and then by the *y* coordinates for each tissue patch. These generate vertical batches that are spatially related and can be passed through the model, as shown by the vertical purity estimates in Figure 5. Since predictions are done on the batch or “bag” level for embedding classifiers, each 120-patch bin in this process generates a single purity prediction.

As shown in Figure 5 (right-most image column), pathologists annotated what they classified as the tumor regions, for several of the WCM slides, to compare to our attention-based areas of interest for the WCM cohort. The regions of high attention weights correspond well with regions within the tissue that pathologists labeled as tumor. In addition, we can infer higher resolution tumor regions for the tumor purity with respect to regions identified using our model. As shown in Figure 5, we have identified specific regions within a tumor region that have low tumor content (as shown by white arrows on the attention map weights). In addition, we compared how the selection of batches may influence the attention maps. We flipped the sorting of tiles for batch generation, to first sort on the *y* coordinates and then the *x* coordinates (horizontal batches). As shown through Supplementary Figure 3, the regions identified do not significantly change based on visual inspection, indicating that our approach is not sensitive to this artificial binning procedure.

We show several different functionalities of our model. First, we show that we can predict the tumor purity, within the entire slide, with a higher accuracy, based on MSE/MAE, as compared to a fully supervised approach, and that we can accurately identify the cancer type within the slide. In addition, we can identify regions within the slide that are tumor positive using attention weights, and we can infer these regions at a higher resolution than what a pathologist would annotate.

## Discussion

In this paper, we present a novel method for determining tumor purity. The method has significant benefits, as it allows for the visualization of tumor regions, as well as regions that have higher vs lower tumor purity. Our method has significant improvement in terms of performance and has added flexibility to allow for various classifications of high vs low tumor purity in a clinical setting depending on biological needs. This model can be applied to many different types of tumors and can be used to identify more pure regions to improve upon (1) patient diagnostics, and (2) improved RNA-sequencing results due to higher tumor purity guided by our methods. In addition, the current approach has a significant amount of flexibility. This approach currently uses pathologist derived purity estimates, but it can easily be applied to other purity measurements potentially using sequencing data if an institution has access to purity scores only for a certain type of purity estimate. Finally, this can be applied for classification between high and low tumor purity for a lot of different thresholds. This is crucial as clinical outcomes based on purity may not be the same between different cancer types. In addition, we can use this in a retrospective study to see how tumor purity using our approach affects the performance of downstream sequencing analyses.

There are some limitations to the current work. The first, is that the distribution of purity scores was skewed. A small portion, 4% of all slides fell below 50% purity, when not considering normal tissue. While this is the inherent nature of the TCGA database, we see this as a potential area to improve model performance in the very low purity regime. Although our WCM test set had examples of low tumor purity slides (Figure 5), a more robust representation could be useful. Our plan could include finding private patient cohorts that have lower purity scores and performing transfer learning or retraining our model on a new set of data, for our model if necessary. Figure 5 shows one example of low tumor purity (row 3), which shows high agreement to the region identified by pathologists, but this example is not sufficient for many conclusions to be made regarding this low purity regime. In addition, there has been work that shows that formalin-fixed paraffin-embedded (FFPE) slides can be used for RNA-sequencing^27^. We can also look to perform transfer learning on a set of FFPE slides, as these slides are used for diagnosis, and these slides maintain a better tissue architecture in most cases compared to frozen sections and are used for diagnostic purposes.

## Methods

### Data Preprocessing

Data was acquired from The Cancer Genome Atlas (TCGA) database, and whole slide images (WSIs) were exported for downstream analysis as svs file format. Slides were taken from six different tumor types. These tumor types were chosen to span the range of tumor purity present in the TCGA database based on previous literature^5^. We analyzed adrenal adenocarcinoma (ACC), lung squamous cell carcinoma and lung adenocarcinoma (LUSC & LUAD respectively), breast invasive carcinoma (BRCA), head and neck squamous cell carcinoma (HNSC), prostate adenocarcinoma (PRAD), and ovarian serous cystadenocarcinoma (OV). We also tested these tissue types to determine Urothelial bladder carcinoma (BLCA) normal tissue slides were used during training, but there was no cancer representation in the training, validation, and test sets to be evaluated (therefore model prediction is for seven classes, although we evaluated metrics for six cancers). Each tumor type was preprocessed separately. Slides were tiled into 512 × 512 patches at 20x resolution (adjusted field of view and downsampling if 40x magnification was used). A color threshold, empirically set for red, green, and blue, was set for each patch to identify the amount of tissue present, and a threshold was set at 40% tissue presence to remove background regions. A Haar wavelet transform was used to filter out tissue that was out of focus due to issues related to the scanner^28,29^. An empirically derived threshold was set for the blur detection to minimize loss of tissue while still removing severely out of focus images (41,632 out of 3,730,063 total patches or ∼1.1%). The slides were split for each tumor such that there was approximately 70% | 15% | 15% for train | validation | test sets for each tissue (Supplementary Table 1). A small fraction of tissues in the training set had less than *N* number of tiles in the slide and were removed from the analysis during training. Slides were split so that each tissue sample was split into one category. Therefore, for cancer samples (−01A labeled through TCGA), all slides from the same patient were grouped into the same split, and for each normal sample (−11A), the slides from a single patient were grouped into the same split. Data distributions as well as predictions for the validation and test sets can be viewed in the supplementary materials (Supplementary Table 2-4).

In addition, we used an WCM independent cohort (outside of TCGA) to validate our model perform. We received 54 de-identified patient frozen section H&E slides from three different cancer types, BRCA, OV, and PRAD, with a pathologist-derived purity estimate to compare with our proposed method.

### Model

We used Pytorch for this study (version 1.1)^30^ and utilized a variant of Resnet34 that is called Resnet34-IBN, which had initially been trained on ImageNet^26,31,32^. The modification to the original Resnet34 model is the incorporation of an InstanceNorm component into the residual blocks (Figure 1C)^33^. In histology images there are noticeable color variations due to differences in tissue preprocessing and stain protocols. InstanceNorm has been used to filter complex color variations (i.e. color shifts or brightness changes), and was utilized for that reason, instead of changing the input of the image through methods such as stain normalization done in other studies^7,26^. To improve model performance and minimize overfitting, we added several different regularization methods. We have added weight decay (L2 Norm) regularization (1×10^−4^), data augmentation (imgaug package) including rotations/flips, coarse dropout, Gaussian noise, hue/saturation/contrast adjustment, and intensity scaling. We use a learning rate 0f 0.005, the stochastic gradient descent (SGD) optimizer, and a bag size of 120 patches, with a batch size of 2 for the MIL set-up. We tested a bag size:batch size ratio of 60:4, 120:2, and 300:1, and the 120:2 gave the best results and therefore were used for all analyses (data not shown). Multi-task learning was performed to analyze both the tissue type present in the slide as well as tumor purity. For tissue type prediction cross-entropy loss was used with inverse frequency of each class used to weight each component in the loss function. For tumor purity the loss used was derived from a previous study, and is used to be able to introduce an ordering to the classification task^34^. Using this ranked loss function, a probability distribution 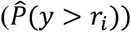, where *r*_*i*_ is the *i*^*th*^threshold used to separate class *i* − 1 from class *i* (where *i* in our case is 8) is generated for each threshold. The additional class, in addition to the 8 that fall above each of the *i* thresholds, is formed for samples that are below all thresholds (i.e. less than 0.09 or less than 9% pure). We binned tumor purity using the following thresholds [0.09, 0.29, 0.39, 0.49, 0.59, 0.69, 0.79,0.89], such that a tumor purity of 0 represents normal tissue (denoted as −11A in the TCGA database) and 1 represents pure tumor. We removed the 0.19 threshold due low representation (14 WSI examples out of 5,860 in the entire dataset or < 0.1% of the total number of slides) within this class. Through this approach, we can set 8 different thresholds when classifying tumors into high vs. low purity, where different thresholds can be used based on biological significance.

In this work, we use an embedding-level MIL classifier. MIL allows for two types of approaches, instance-level, where classification can be done on each individual component associated with a bag, and embedding-level, where the bag is classified but the meaning of each individual instance is lost. We chose to use the embedding level approach as previous literature has shown that the attention mechanism allows for importance ranking of individual instances within a bag^20^. In traditional multiple instance learning, under the collective assumption, all training instances in a given bag contribute equally to the final prediction^19^. We modify this, based on previous works, to include a weighted average of instances, which is learned through an attention mechanism. We use the gated attention mechanism proposed previously^20,35^, where the weights derived for each patch *k* is calculated by:

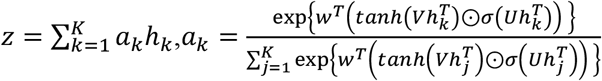

In these equations, *h*_*k*_ represents the specific feature embedding, *V, U* ∈ ℝ^*LxD*^ are the same dimension where *L* has been set to 128, and *D* is the size of the feature embedding, and *w* ∈ ℝ^*Lx*1^ to generate the weights, which then go through a softmax function which is used to pool feature embeddings^20,35^. The idea of an MIL-embedding classifier is that we get a prediction for a set of patches, but do not have to predict each individual patch. We use the attention weights to identify different regions in the tissue that have different weights for prediction.

The final loss function was represented as *L* = *L*_*purity*_ + α*L*_*tissue*_, where α is set to 0.125 to allow for an equal influence for both prediction tasks based on empirical testing. The model was trained using a single GeForce GTX 1080 GPU for 9 epochs and we chose the model with the lowest validation loss. Supplementary Figure 1 shows the training loss and validation loss as a function of training epochs.

### Dataset Generation

The slides in the dataset can consist of multiple individual histological slices of the same tissue. To tackle this problem, these slices were grouped into geographical subregions. The subregions were later used to train, validate, and test the predictive models. First, we generated a slide position matrix, consisting of the x and y coordinates of all the tiles per slide. Next, to identify the individual tissue slices, we used density-based spatial clustering of applications with noise (DBSCAN) over the plotted data^36^. We used ϵ of 0.3, where ϵ is the maximum distance between two samples for them to be considered in the same neighborhood^36^. The clusters identified by the algorithm represent the individual tissue slices in a slide. Finally, the tiles in each cluster were sorted first by *x* and second by *y* and were split into subsets of sizes 120 (to prevent memory constraint issues when running the deep learning models). The resulting subsets were used to train, validate, and test the predictive models.

### Data Analysis and Visualization

To visualize the features developed from the MIL-embedding, we used t-distributed stochastic neighbor embedding (tSNE) to reduce the dimensionality of feature vectors extracted from the model. We performed dimensionality reduction on both arms of the multi-task learning model, right before the last linear layer, and visualized the purity score and tissue type label by color for each bag based on the slide label. In addition, tissue patches were reconstructed with the predicted label to generate heatmaps for prediction and attention weights (normalized to span from 0—1) were used to identify different regions on the slide that were weighted differently by the model. We multiplied the attention weights for the tumor type prediction by the predicted tumor purity, to better identify the tumor regions of interest.

As mentioned, the loss function for purity score prediction generates a probability vector ŷ, which is of length *K-1*, where *K* is the number of different categories. We generate ROC curves, using the package scikit-learn, based on these probabilities for each of the thresholds set. We average the probability vectors for each slide to get a final probability vector that we use to get the ROC curve and subsequent AUC value. For each cancer type, we calculated accuracy 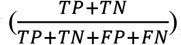, precision 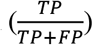, and recall 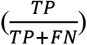 based on the number of true positive (*TP*), true negative (*TN*), false positive (*FP*), and false negative (*FN*) tissue type predictions generated by the model (selecting the tissue type with the highest probability). We aggregate the predictions from all tiles the entire slide and do a majority voting to get slide level tissue type predictions.

We compared the tumor purity score predictions by our model to the predictions generated in a study by Fu et al.^25^. In this paper, the authors use label smoothing to predict patches for each tile in a fully-supervised approach, where the true label is given as the tumor purity for the sample. For the comparison, we calculate MSE and MAE for both studies based on true versus predicted tumor purity scores of 701 slides that were present in the test datasets of both studies (Table 3). Our model generates a list of probabilities per tile, where the i^th^ element of the list corresponds to the probability the tile has a tumor purity at least within the i^th^ bin. To calculate a tumor purity score of each tile, we used a probability cutoff of 0.5. Next, we calculated the average tumor purity score of all tiles in a slide to get a tumor purity score on a slide level. Finally, to compare the predictions generated in two studies, we used the MSE and MAE functions from the sklearn.metrics module.

Plots were created in python using matplotlib and seaborn.

Code for training and the trained models are publicly available on GitHub at https://github.com/ih-lab/wsPurity

## Contributions

M.B., P.K., I.H and O.E. conceived of the project. M.B. and V.G. carried out experiments and wrote the manuscript draft. MS, AS, TK, MA and JMM assisted with acquiring and the interpretation of the internal EIPM cohort. MA and JMM provided histopathology interpretation. I.H. supervised the study. All authors read, edited, and approved the final manuscript.

## Acknowledgments

This work was supported by start up funds (Weill Cornell Medicine) to I.H. The results shown here are in whole or part based upon data generated by the TCGA Research Network: https://www.cancer.gov/tcga.

## Supplementary Material

**Supplementary Figure 1:**
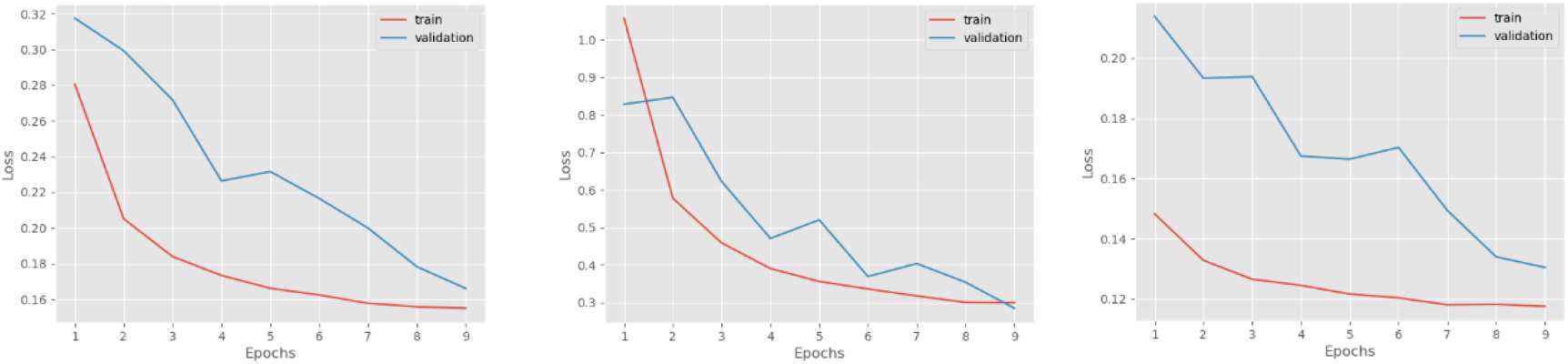
Train and Validation Loss for (1) Overall Loss, (2) Tissue Type Classification Loss, (3) Purity Classification Loss for the 10-epoch training process.

**Supplementary Table 1.** Distribution of samples per cancer/tissue type.

| Cancer Type | Tissue Type | Training | Validation | Testing |
| --- | --- | --- | --- | --- |
| ACC | Cancer | 68 | 16 | 6 |
|  | Normal | 2 | 1 | 1 |
| BRCA | Cancer | 966 | 232 | 204 |
|  | Normal | 230 | 68 | 57 |
| HNSC | Cancer | 469 | 100 | 88 |
|  | Normal | 26 | 8 | 6 |
| LUAD and LUSC | Cancer | 1016 | 185 | 209 |
|  | Normal | 388 | 114 | 84 |
| OV | Cancer | 329 | 71 | 83 |
|  | Normal | 109 | 27 | 26 |
| PRAD | Cancer | 382 | 77 | 84 |
|  | Normal | 78 | 22 | 18 |
| Total | - | 4063 | 921 | 866 |

**Supplementary Figure 2:**
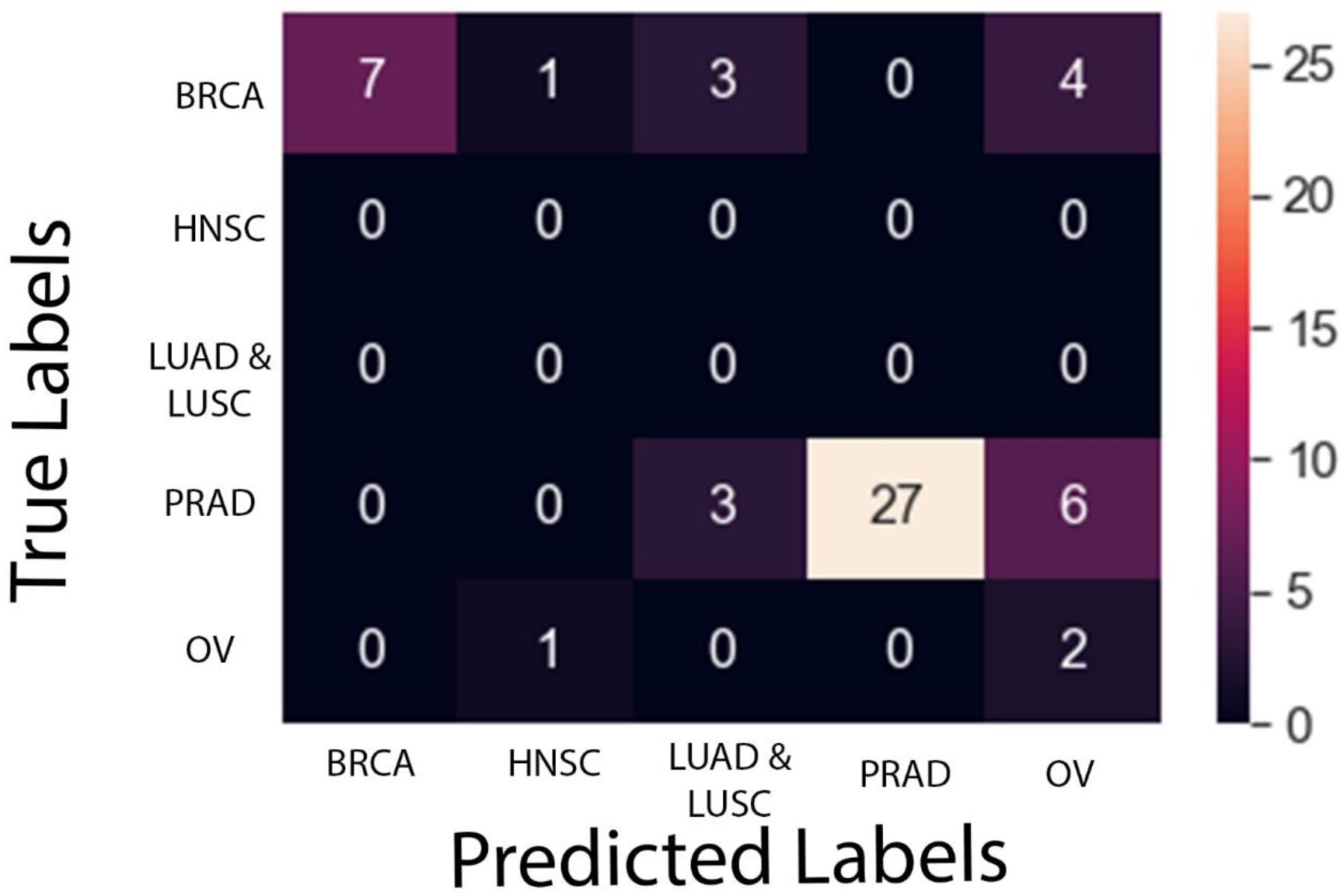
Confusion matrix for tissue type prediction on the WCM cohort of patients

**Supplementary Figure 3:**
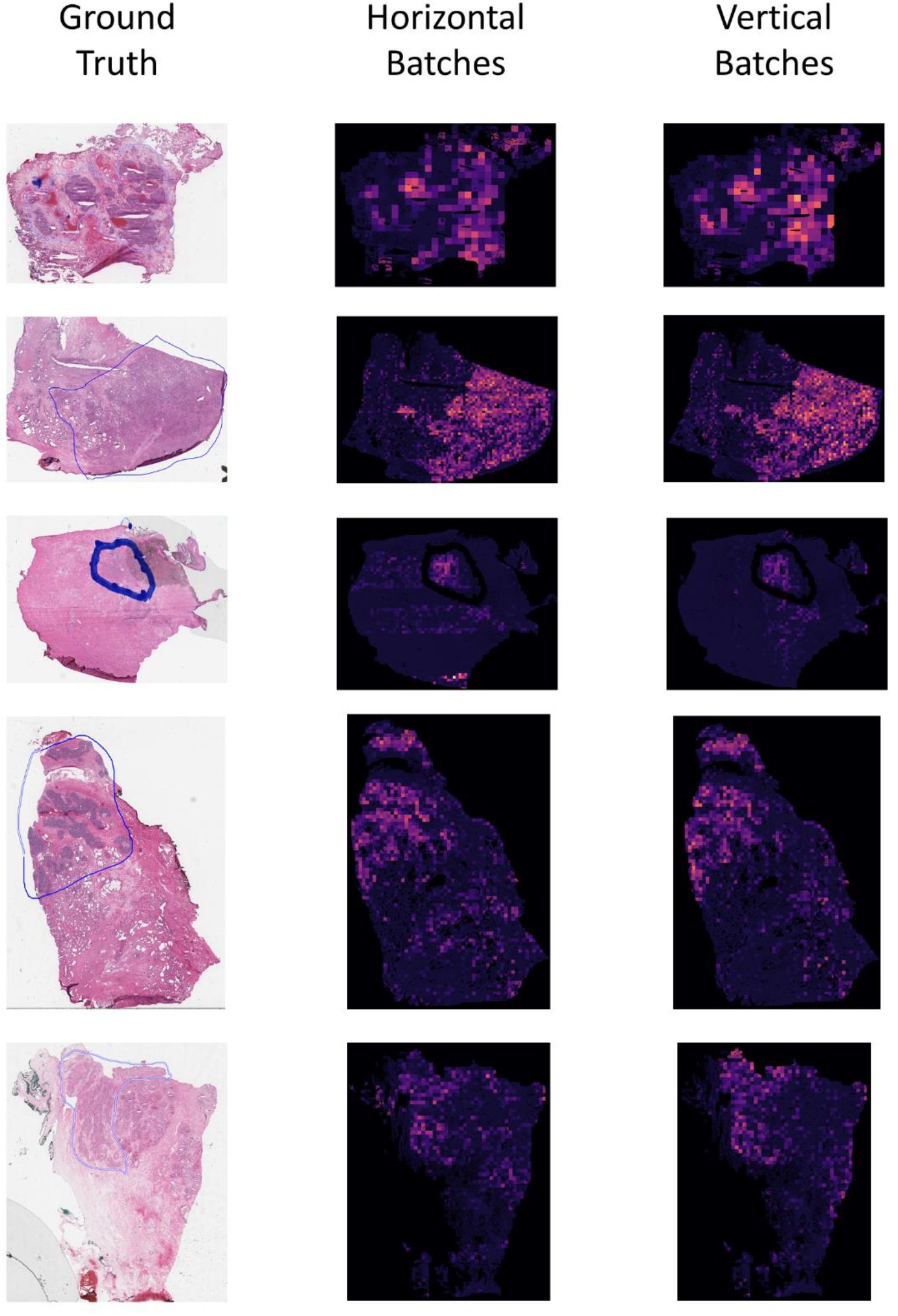
Comparison of tumor attention maps using batches derived from tiles sorted using *x* then *y* coordinates (vertical batches) as well batches derived from tiles sorted using *y* then *x* coordinates (horizontal batches)

Supplementary Table 2-4: (2) Data splits for training, validation and test set for TCGA cohort. (3) Predicted values on the validation set for TCGA cohort. (4) Predicted values on test set for TCGA

